# Nanopore Translocation of Topologically Linked DNA Catenanes

**DOI:** 10.1101/2022.08.10.503522

**Authors:** Sierra N. Rheaume, Alexander R. Klotz

**Affiliations:** School of Physics, Georgia Institute of Technology; Department of Physics and Astronomy, California State University, Long Beach

## Abstract

The electrical signal associated with a biopolymer translocating through a nanoscale pore depends depends on the size, topology, and configuration of each molecule. Building upon recent interest in using solid-state nanopores for studying the topology of knotted and supercoiled DNA, we present the first experimental observations of topologically linked catenanes translocating through a solid-state nanopore. Using restriction enzymes, linked circular molecules were isolated from the mitochondrial DNA of *Crithidia fasciculata*, a structure known as a kinetoplast that is comprised of thousands of topologically interlocked minicircles. Digested kinetoplasts produce a spectrum of catenane topologies, which are identified from their nanopore translocation signals by spikes in the blockade current associated with the topological linkages. We identify the translocation signatures of 2-catenanes, linear and triangular 3-catenanes, and several types of 4- and 5-catenanes as well as more complex structures. Measurements of the translocation time of 2- and 3-catenanes suggest that topological friction between the linkages and the pore slows the translocation time of these structures, as predicted in recent simulations.

## Introduction

Nanopore sensors detect the electrical current associated with the passage of an ionic fluid through a nanoscore pore in a membrane. Biomolecules translocating through the pore cause current blockades that may be used to detect or characterize the molecules.^1^ Nanopore sequencing, in which the genetic sequence is reconstructed by using the current blockade associated with individual nucleotides, has recently become a viable tool for long-read-length sequencing using biological pores.^2^ Solid state nanopores offer greater versatility and tunability than biological pores, and have been used for identifying the binding of individual proteins bound to DNA^3^ and for direct protein detection. ^4^ The development of nanopore technology has grown in step with an improved understanding of polymer dynamics during the translocation process.^5,6^ Recently, there has been theoretical interest in the physics and rheology of topologically complex polymers, including rings, ^7^ branched chains,^8^ knots,^9^ and catenanes. ^10^ Many of the experimental investigations have used DNA as a model system. Recently, experiments examining knotted DNA in solid state nanopores have provided information about the frequency of stochastic knots in DNA,^11^ the size distribution of knots in DNA,^12^ and the mechanisms by which knots deform as they translocate.^13^ While knots in linear strands are transient and can untie before or during translocation, linked-ring catenanes are topologically robust. Experiments with catenanes have the potential to explore so-called “topological friction” in a way that previous experiments have not been able to. While there have been simulation studies of the catenane translocation process, ^14,15^ experiments have been lacking.

Much of the interest in topologically complex translocation concerns topological friction, a phenomena whereby the altered topology of a molecule influences its tribological interactions with its environment. Hypothesized to explain the ejection of knotted genomes from virus capsids, ^16^ topological friction has been studied in the context of knot diffusivity^17^ as well as the translocation of knotted DNA molecules through pores.^18–20^ Experiments with knotted translocation have provided only transient data for topological friction.^11^ In contrast, simulations of catenane translocation predict much stronger topological friction effects, including a jamming associated with the linkage between catenaned rings that has a significant influence on the translocation time.^14,15^

Kinetoplast DNA (kDNA), often described as “molecular chainmail,” is a complex DNA structure consisting of several thousand linked circular molecules around 2500 base-pairs in length, known as minicircles, and several dozen larger circular molecules called maxicircles, around 40,000 base-pairs in length.^21^ kDNA is found in the mitochondria of Trypanosome parasites, which are responsible for diseases such as Leishmaniasis, Chagas Disease, and Sleeping Sickness. kDNA is part of a complex gene-editing apparatus that allows metabolic proteins to be expressed by modifying the mitochondrial messenger RNA.^22^ Kinetoplast biology is an active area of parasitology research^23^ and kDNA has recently been explored as a model system for two-dimensional polymer physics. ^24,25^ Here, we use kinetoplasts from *Crithidia fasciculata* as a source of topological linkages to investigate catenane translocation. Restriction enzymes can linearize minicircles within kinetoplast networks, which can release catenanes of various complexity into the solution (Fig. 1a). ^26^ These released catenanes make up an ensemble of topologically diverse molecules that we can observe using a solid state nanopore (Fig. 1b).

**Figure 1:**
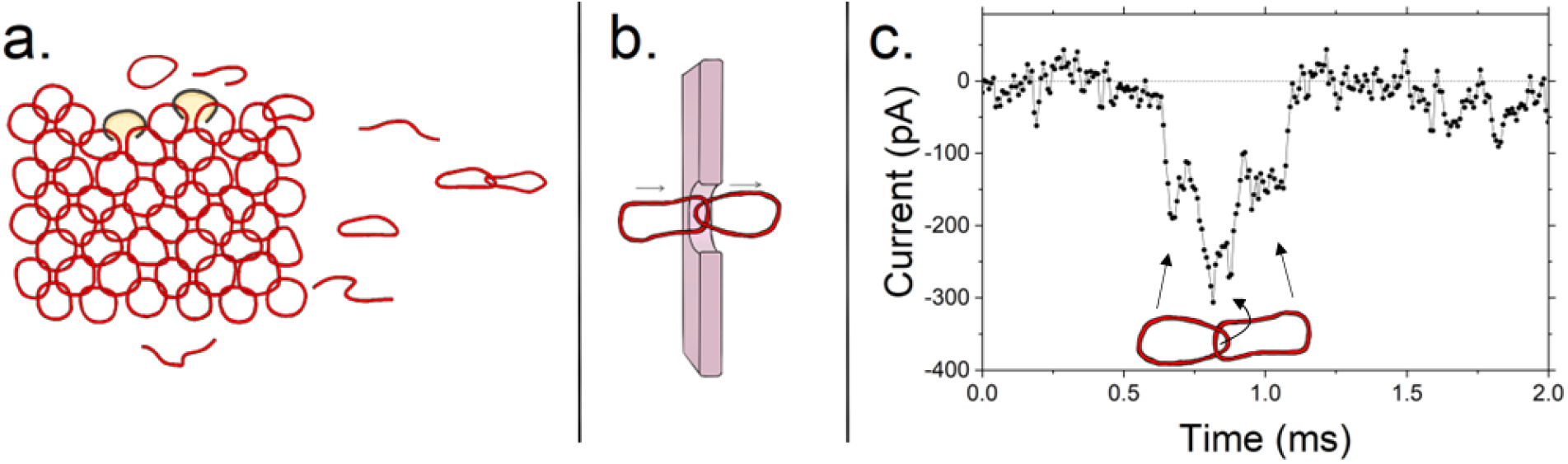
a. Schematic of kinetoplast DNA (red) being digested by XhoI restriction enzymes (yellow/black) which linearize molecules and can release isolated circles and catenanes from the network. b. Schematic of a Hopf link or 2-catenane translocating through a pore. c. The translocation current from an experiment showing two baseline drops connected by a deeper blockade, which we interpret as the linkage between two Hopf-linked minicircles.

## Experiments

Kinetoplast DNA from *Crithidia fasciculata* (TopoGEN) was digested with XhoI restriction enzyme (schematic in Fig. 1a) in CutSmart buffer (New England BioLabs) in a 50 *µ*L aqueous solution containing 1 *µ*g of kDNA, 5 *µ*L of CutSmart buffer and 1 *µ*L of XhoI at the supplier’s stock solution. The solution was placed in a 37^*◦*^C water bath for up to 20 minutes. Proteinase K was added to the solution after the allotted time to suppress additional digestion. Following the results of Chen et al., ^26^ this produced a spectrum of topologically connected DNA molecules with a frequency that depends on the degree of enzyme digestion and the underlying connectivity of the kinetoplast network. The digested DNA solution was diluted in 4 M lithium chloride solution to approximately 1 nM.

Solid state nanopores in silicon nitride membranes embedded in microfluidic *cis* and *trans* channels were enlarged using controlled dielectric breakdown^27^ by filling the channels with 4M LiCl and applying 10 volts across the membrane. The typical size of the nanopores is between 30 and 50 nm. The NanoCounter (Ontera, Inc.) was used both to enlarge the nanopores and to facilitate and detect the translocation of DNA by applying a 100 mV potential difference across the pore and measuring the current through it (schematic in Fig. 1b). Thousands of DNA translocation events were observed in each experiment, many of them containing short spikes of excess current blockage in the interior of the molecule. These spikes are similar to those observed in knotted molecules, reported in previous studies. ^11–13^ A characteristic molecular translocation signal associated with a Hopf link (2-catenane) can be seen in Fig. 1c.

In order to determine the topology of translocating molecules, we predicted the translocation signal of each expected configuration with up to five catenaned links, ^26^ and used the SPIRaL classification scheme developed by Sharma et al. ^12^ to predict their translocation signal based on the number strands expected to occupy the pore as the molecule translocates. For example, a Hopf link is expected to have two strands in the pore as the first minicircle translocates, four as the linkage translocates, and two as the second translocates, and would have a SPIRaL code of 2-4-2, which is observed in Fig. 1c. Similar to previous work used to classify knots, we examined the translocation signal of molecules that were localized above the median translocation time and mean current blockade, and classified them into their expected topologies. Fig. 2 shows four of the simplest topologies, and a listing of all observed topologies, their SPIRaL classifications, predicted signals, and observed signals, can be seen in SI Figure 1, with unobserved topologies listed in SI Figure 2. We observed the 2-catenane, both forms of the 3-catenane, three of five 4-catenanes, and six to seven of eleven 5-catenanes, as well as structures consistent with maxicircles linked to minicircles.

**Figure 2:**
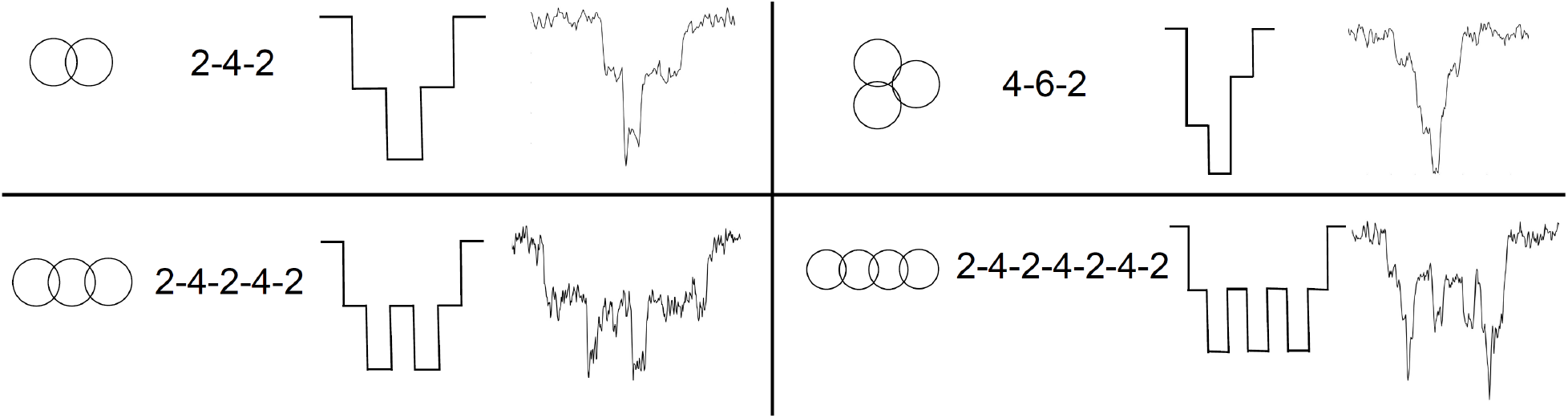
Four of the simplest expected catenane topologies (2-catenanes, linear and triangular 3-catenanes, and the linear 4-catenane). Shown are a diagram of their topology, their predicted SPIRaL classification based on the number of concurrent DNA strands within the pore during translocation, a predicted translocation signal based on the SPIRaL classification, and an example of an observed signal.

## Results and Discussion

Translocation statistics are visualized on a mean current blockade-vs-translocation time scatter plot. A plot from a single experiment can be seen in Fig. 3a, where the majority of translocations lie in a banana-shaped region of constant charge deficit, associated with single minicircles. It is not possible to distinguish folded linear molecules with unfolded circular molecules, thus this region contains the aggregate of both open and closed minicircles. We observe a secondary region at slightly longer time which we associate with fragments from cleaved maxicircles. Compared to experiments with standard linear or circular molecules, we observe a significant population of translocations away from the banana. Observing these translocations reveals features associated with molecular catenanes, which we have examined, classified, and color-coded by their topology in Figs. 3a and 3b.

**Figure 3:**
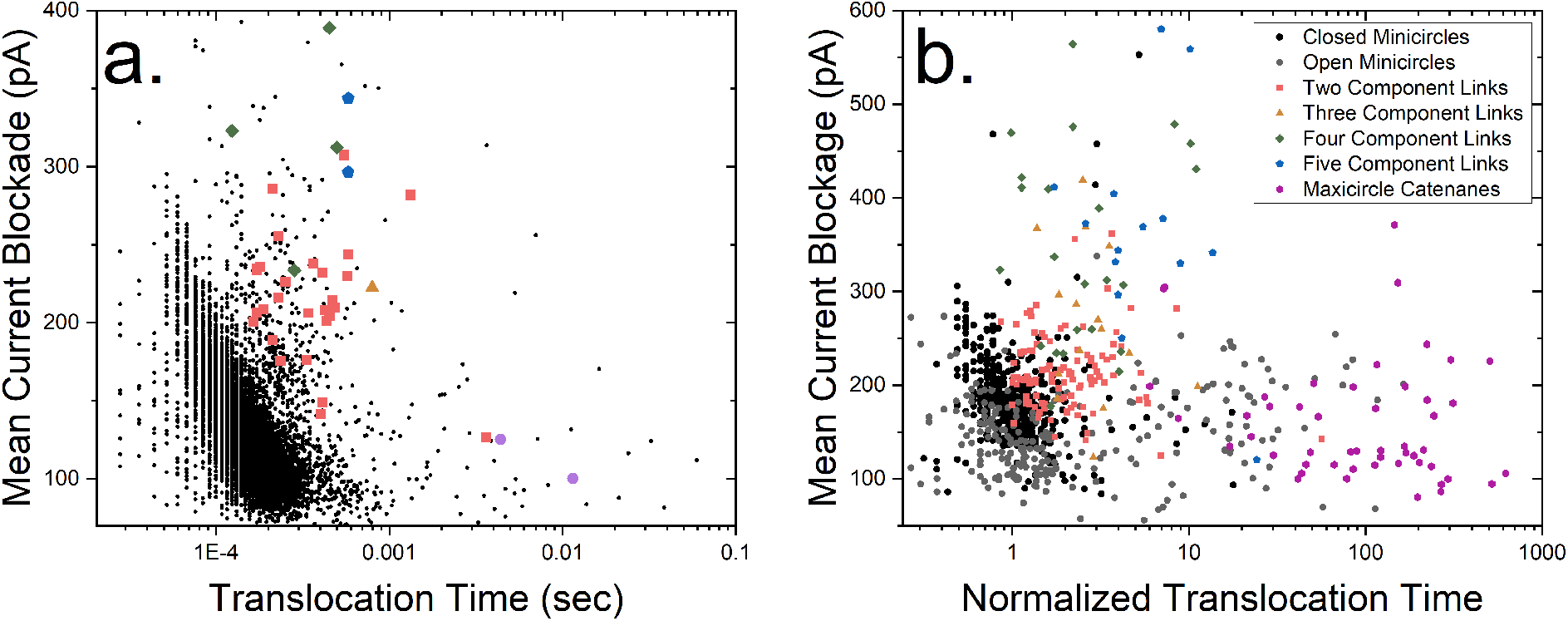
a. Current-time translocation scatter plot from a single experiment showing the open and closed minicircles in black and more complex topologies colored according to the legend in (b). The secondary cluster to the right of the main minicircle cluster likely contains cleaved maxicircle fragments. b. Collected current-time scatter plot showing nontrivial minicircle topologies aggregated over several experiments, color-coded according to the number of minicircles in the translocating structure. The grey and black data are from experiments with isolated linear and circular minicircles. Translocation time data are normalized to the peak time of the (linear+circular) minicircle histogram.

We are interested in the mean translocation time of exotic topologies, which is pore size-dependent. The Ontera NanoCounter creates pores through controlled dielectric breakdown, and produces a stochastic range of pore sizes compared to those created by electron or ion beam milling. In order to normalize experimental data recorded on different devices, we examine a histogram of linear and circular minicircle translocations for each experiment, find the peak to define a typical minicircle translocation time, and divide each exotic translocation time by the peak time for that experiment. Because the single-minicircle data is an aggregate of both linear and circular molecules, the peak time is in between the peak time of linear and circular molecules, and the measured ratio of exotic topologies should be considered a lower bound on the ratio of each topology to a single minicircle.

An aggregated scatter plot in Fig. 3b shows normalized data from conventional (linear and circular) and exotic topologies. We are able to distinguish between open and closed minicircles in this plot by performing experiments with isolated linearized and decatenaned minicircles. The data in Figure 3b show the normalized translocation times and mean currents of exotic translocations color-coded according to their ascertained topology. Most observed catenanes were Hopf links, followed by linear and triangular 3-catenanes. We have observed a significant population of combined maxicircle-minircle complexes. We have observed nearly every expected topological structure of up to five minicircles at least once (but not in sufficient numbers for meaningful statistics). Generally speaking, we observed significantly fewer catenanes than predicted by the analysis of Chen et al. ^26^ The intensity of a gel band is proportional to the mass of the DNA in the band rather than the number of distinct molecules; thus the larger catenanes are easier to detect compared to a nanopore experiment. In addition there may be differences in the nanopore capture process due to topology and as such, we refrain from making inferences based on the observed topological distribution.

The normalized mean translocation times of the three most common exotic topologies are found in Table 1. As a point of comparison we show similar data from a simulation study ^14,15^ using the unknot translocation time as the basis of normalization. As the simulations use a smaller pore size and flexible (not semiflexible) chains, comparisons should be treated as qualitative. Due to being observed in a much greater quantity, we regard our 2-catenane results as being the most significant.

**Table 1:** Ratios of the mean translocation time for 2-catenanes, linear 3-catenanes, and closed (triangular) 3-catenanes to the peak translocation time of linear and open minicircles, with standard error. Computational data from Carraglio and Orlandini ^15^ is provided as a comparison. Because the nanopore cannot distinguish between linear and closed minicircles, the measured ratios should be treated as a lower bound.

| Topology | Exp. Time Ratio | Comp. Time Ratio<sup>15</sup> |
| --- | --- | --- |
| 2-Catenane | $2.7 \pm 0.5$ | 2.7 |
| 3-Catenane | $3.9 \pm 1.1$ | 4.8 |
| Triangle | $2.5 \pm 0.3$ | 4.0 |

Arguments may be made that topological complexity can either speed up or slow down a molecule in tight confinement: a speed-up effect due to the effective shortening of the molecule due to a knot, or a slowing effect predicted to arise from topological friction. If topological friction dominates over shortening, we expect the 2-catenane to take more than twice as long to translocate as an isolated minicircle. Our observed ratio of 2.7*±*0.5 indicates that topological friction is significant. The 3-catenane is expected to take more than 1.5 times that of the 2-catenane to translocate if friction dominates, although our data are not precise enough to make that observation. The shorter translocation of triangles compared to linear 3-catenanes indicates that shortening can also be a significant effect, similar to the shorter translocation times for knotted DNA seen in previous experiments. ^11,12^ Another comparison may be made between Hopf links and single loops with twice the countour length, although direct comparison is difficult because of the superlinear translocation time exponent. ^28,29^ Hopf links are predicted to translocate slightly faster than double-sized single links^15^ due to a balance of friction, shortening, and length-dependence effects, a phenomenon that can be clarified in a future experiment.

One of the predictions of Caraglio and Orlandini is that topological friction causes the linkage between molecules to dwell in the pore for long periods of time.^14,15^ This would manifest itself experimentally as an extended drop in the measured current. Fig. 4a shows several 2-catenane translocations with a broad distribution of linkage spike widths is observed. Many are short, but several last for a time comparable to the translocation of a complete minicircle, suggesting that jamming effects are present. We measured the linkage translocation time by counting the number of contiguous time points in each molecule the translocation signal was below a threshold of 1.5 times the median current blockade level. A histogram of the linkage translocation time is shown in Fig. 4b, with a peak at short times but a broad tail with a few observed long outliers. This lends further support to our assertion that the linkage between two linked minicircles experiences topological friction within the pore.

**Figure 4:**
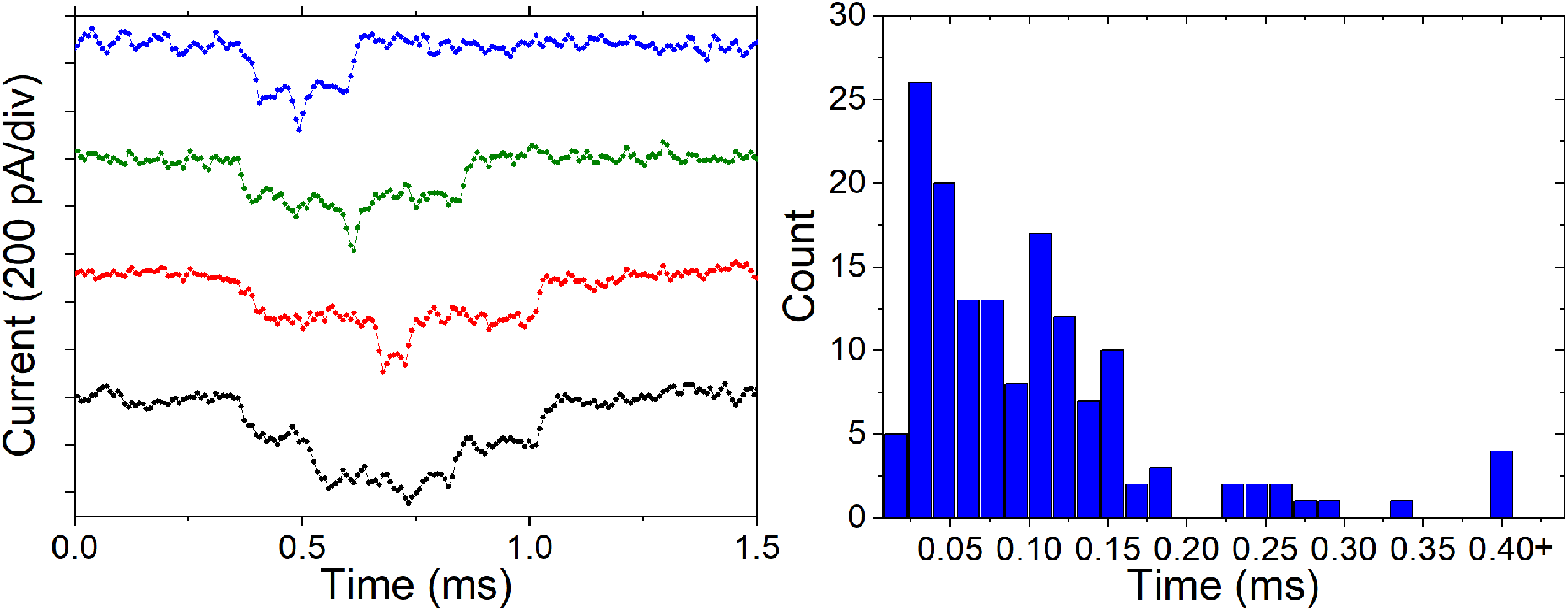
a. Several current traces of 2-catenanes or molecular Hopf links. Variation can be seen in the width of the central spike. b. Histogram of observed spike distribution times for the population of translocating 2-catenanes.

## Conclusion

We used a solid-state nanopore to study the translocation signature of DNA catenanes, isolated from kinetoplasts using restriction enzymes. We observe a rich spectrum of topological structures as determined by the spikes in their transocation signal. The topology-dependence of the translocation time of these catenanes indicates the presence of topological friction between the linkages and the pore, as predicted by simulations. This study used pores that were up to 12 times larger in diameter than the minimum width of circular DNA: future experiments may probe translocation in sub-10 nm pores comparable to the minimum width of a circular DNA molecules, and use topoisomerase decatenation rather than restriction digestion to produce a broader spectrum of catenaned topologies. Many simulation studies focus on increasingly exotic topological linkages between pairs of rings, but we hope this work inspires investigation into the translocation of more complex Hopf-linked rings, ^30^ similar to those now observable experimentally.

