## Supplementary Information for "Nanopore Translocation of Topologically Linked DNA Catenanes"

August 10, 2022

### Classification of Observed Topologies

In their paper on kinetoplast topology, Chen et al. [1] report SEM observation of various catenane structures and use that to refine their estimates. Here in Figure 1 we report the zoo of observed topologies, as well as one structure consistent with a catenane of maxicircles and several minicircles.

For each topology predicted from combinatorics, we calculate a "SPIRaL" classification [2] based on the number of concurrent strands within the pore as the molecule translocates, then draw a barcode based on that numeric classification, and compare non-bananular translocation signals to these codes to ascertain molecular topology. We note that we generated the predicted barcodes before undertaking analysis of the data. The final trace shown is likely a structure consisting of a maxicircle linked to several minicircles.

Several of the more complex structures predicted by Chen et al. [1] were not observed. They are shown in Figure 2.

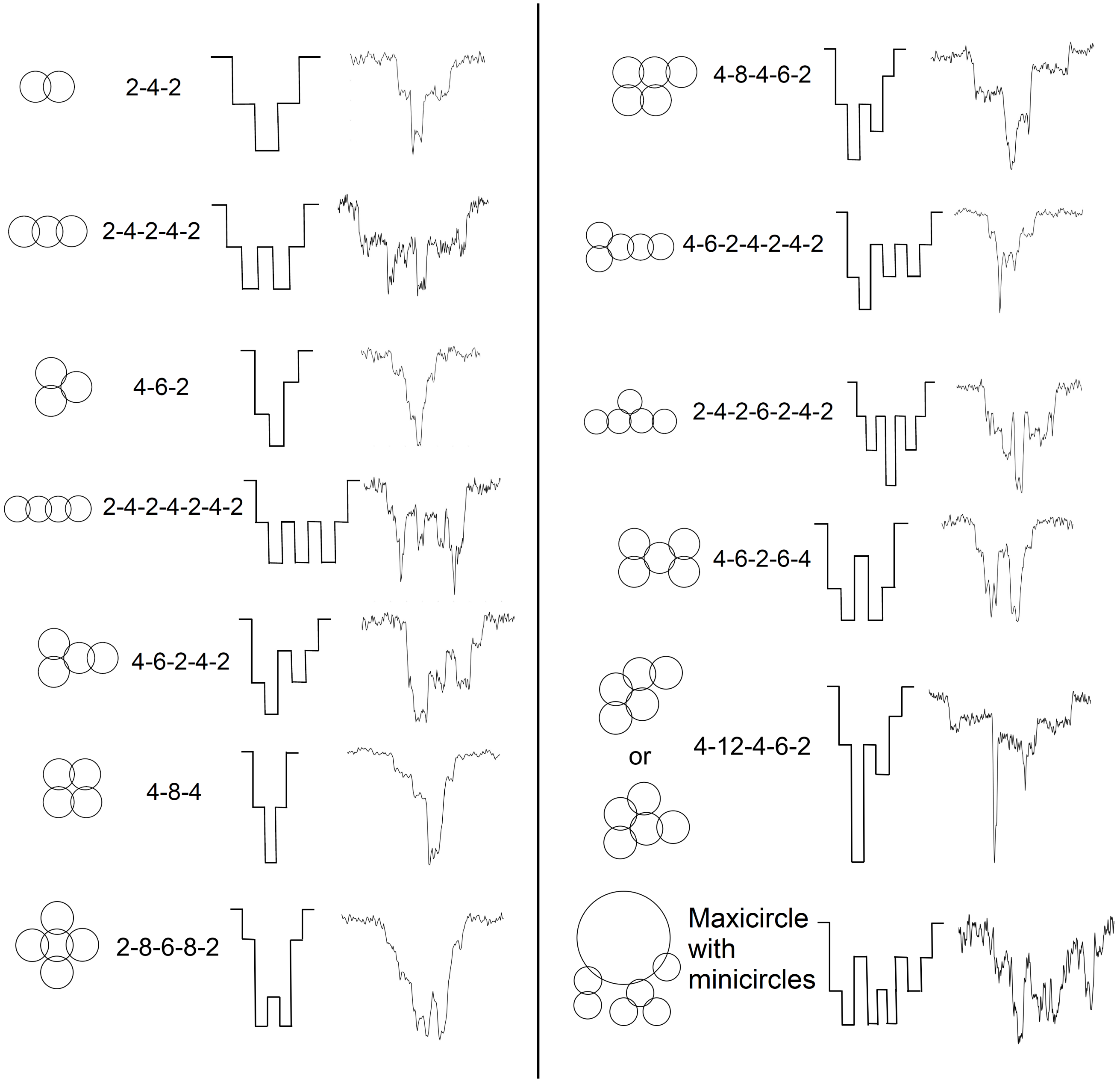


Figure 1: Table of linked topologies, their SPIRaL classification [2], a schematic of a translocation, and an observed translocation. The final topology likely includes a maxicircle linked to several minicircles, with unknown exact topology.

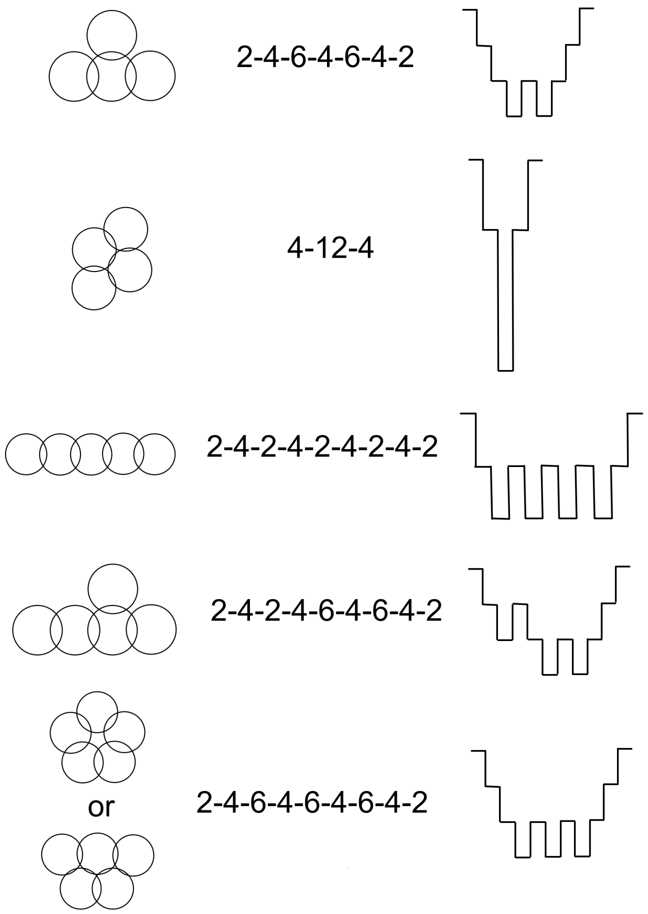


Figure 2: Catenated structures, their SPIRaL classification, and predicted translocation structure, for topologies that were not observed.
